# Defocused reflectance imaging for low numerical aperture, large field of view quantitative live cell imaging studies

**DOI:** 10.1101/2025.03.24.645046

**Authors:** Sophie Bulloch, Tienan Xu, David Herrman, Paul Timpson, Tri Giang Phan, Yu-Hsuan Lin, Makoto Banno, Yean Jin Lim, Woei Ming Lee

## Abstract

High-throughput live-cell imaging within incubator environments often necessitates a compromise between optical resolution and instrument/computational complexity. In this work, we demonstrate that controlled defocusing, a default function in all optical microscopes, can be utilized as a primary contrast mechanism for large-scale cell analysis (7000 cells per field of view). Our results suggest that even at a low numerical aperture (NA ∼ 0.01) and using just epi-illumination, defocused images can generate a uniform negative contrast across the cell body. This negative contrast improves automated cell segmentation efficiency over a wide field of view compared to in-focus imaging. We further evaluated the accessibility of this defocus imaging approach for both 2D and 3D cultures by implementing an automated cell segmentation protocol on a commercial off-the-shelf digital Universal Serial Bus (USB) microscope. The compact form factor of the digital USB microscope facilitates minimal pixel sampling, enabling high-throughput single-cell detection and continuous tracking across a large adherent cell population over several days. Our assessments of the utility of defocus imaging for 2D and 3D tissue cultures were further supported by monitoring the negative contrast changes during the migration and dissociation of 3D tissue spheroids. Our results show that negative contrast profiles from defocus images enable the quantification of cell proliferation, division, migration, and cell-to-cluster dissociation within standard culture environments. This defocusing methodology offers a scalable approach to extensive high-content screening through simplified instrumentation controls.

**Key points:**

- Defocused images of adherent cells using low NA optics homogenizes intracellular intensity hotspots, creating uniform negative contrast for whole cell segmentation; similar to shadow imaging in fluorescence.
- Inverting look-up-table (LUT) combined with defocus imaging on large population of adherent cells in standard culture flask taken by standard digital USB microscopes allow for rapid, routine tracking of thousands of cells per field of view.
- Negative contrast image profiles from defocusing enable the quantification cell proliferation, division, migration for 2D monolayer and 3D tissue spheroid cultures.

---

Osterberg and Smith [1] reported that in telescopic imaging that while defocusing increases the Airy disc (single point spread function), the increased side lobes of Airy disc in fact improves the ability to resolve adjacent points i.e. Rayleigh criterion. In biological imaging, the use of defocused images can improve imaging contrast because the wavefront shift incurred when collimated light passes through organelles creates gradients of intensity at the imaging sensor. These intensity hotspots were shown to enable tracking membrane fluctuations and vesicle movements [2-5]. Another use of defocussed images of whole cells have shown to achieve quantitative tomography imaging of live cells [6]. Yet, the direct deployment of defocus imaging to improve image contrast remains underutilized in live-cell imaging due to critical questions over its performance. Some of these questions include; (1) How defocused images of cells affect modern cellular image segmentation workflows in 2D and 3D cell cultures? (2) What quantitative benefits would defocus imaging bring? (3) Does defocus imaging operate in the same way in reflectance-only imaging mode since previously defocus imaging where all demonstrated in transmitted-only imaging modes?

In this paper, we set out to address the above questions with a primary focus to determine if defocus imaging can be used for routine 2D and 3D cell culture studies. We chose a condenser-free reflectance imaging setup to minimize physical integration challenges. With the condenser, the reflectance microscopes can easily fit into any incubator (≤ 30 litres). Three distinct condenser-free reflectance modes, consisting of scattering, oblique, and epi-like illumination, were evaluated. To measure segmentation performance across the three different reflectance modes, we used a Dice score to quantify the overlapping masks of cells grown in regular culture flasks [7]. For practical implementation of defocus imaging, we proceed to utilise a consumer-grade digital USB microscope that can operates in a reflectance mode and validate defocused imaging modes for quantitative tracking and measurement of single cell movements [8-11] in long term 2D cell cultures. This allows us to investigate if defocussed images can operate with sufficient throughput and resolution to cater to long-term tracking of large cell populations. Finally, we then explored how defocus imaging fare when used to measure morphological changes in 3D thick spheroids. In summary, this study provides a comprehensive and practical evaluation as to whether defocus imaging is suitable for day-to-day 2D and 3D cell culture studies and the approaches to use defocus images for quantifying general cell culture dynamics.

## 1. Results

### Is there an optimal defocusing illumination to achieve negative contrast?

In this section, we determine the optimal level of defocus, the illumination type for most effective image segmentation across various cell culturewares. We compared defocused images from each reflectance mode to determine which angled illumination is better suited for image thresholding against changes in background intensities. Dice score [12] was then used to determine segmentation efficiency using manual segmented image as ground truth to determine overlapping regions. Figure 1 (i) illustrates the experimental setup, featuring a multimode fiber (NA = 0.22) coupled to a white light-emitting diode (LED). This configuration uses a separate illumination geometry and avoids the use of a beam splitter as that it can facilitate broad adoption across imaging systems that do not necessarily support integrated light paths i.e. use of dichroic and image splitting mirrors. Independent illumination systems have been used in lightsheet imaging [13, 14] which demonstrated flexibility across different reflectance imaging system. We showed in Fig. 1(ii) that brightfield images of single and aggregated murine cells (L929, ATCC® CCL-1™) display difference in intensity gradient and background intensities. In particular, in scattering mode, the images displays a dark background with uneven light distribution across cells and spheroids, akin to darkfield imaging. Oblique and epi-illumination provide a uniform background; however, oblique illumination results in an uneven grayscale, while epi-illumination reveals distinctive intensity changes within the cell body under different defocusing distances. Cells were cultured in methylcellulose (MC) media after a three-day grown in T75 Flask (Nuc™, Thermofisher) captured at various defocus distances (-5 µm, 0 µm and +5 µm) respectively. We then applied a series of post-processing steps to retrieve the image masks from the three different illumination modes. The post-processing steps (background subtraction, thresholding, and binary filtering) illustrated in Figure 1 (iii) (left) were undertaken to retrieve the image masks (red outlines). Examples of the results are shown in Figure 1 (iii) (right). Figure. 1 (iv) all the images and the respective outline (red line) overlayed with the ground truth (manual segmented). The overlayed masks and the raw images were then used to determine a series of Dice score. The Dice score [12, 15] were used to measure mask similarity, where 1 represents the perfect outline for image segmentation. As shown in Fig. 1 (v), the qualitative comparison of segmentation efficiency using the Dice score indicated that the score is highest for defocusing under the regular epi-illumination mode. Whilst the overall Dice score for different epi-illumination (Dice < 0.13) are much lower than fluorescence approaches (dice >0.8) [12], it is evident that the both defocusing mode has the higher segmentation efficiency according to the Dice score than others modes except for oblique illumination[16]. This findings suggest that at low NA objectives, defocus images functions similarly to shadow imaging where an inverted look up table (LUT) [17] increases image contrast that improve segmentation efficiency Qualitatively, the images shown in Figure 1 (iv) indicate that in-focus and out-of-focus images taken from scattering and oblique illumination generate high-intensity hotspots that affect the thresholding process when identifying boundaries of each cell, despite having a high visual contrast of the cell edges and organelles. However, defocused images for epi-illumination stand in contrast to in-focus images. Internal intensity spots within cells, which exhibit higher intensity variance that fails to effectively highlight the whole cell body, were blurred out in the positively defocused image. As a result, a positive defocused image creates a uniform area of low intensity that effectively blurs any intensity hotspots, thus providing an ideal contrast change for segmentation [6]. Whilst the dice comparison shown in Fig. 1 (v) indicated that both defocused images (dice = 0.113, and 0.1107) have close to oblique (dice = 0.1362). At closer examination to compared between Figure. 1 iii), the positive defocusing creates a uniform contrast across the cell body where internal cellular structures is observed in oblique illumination introduce noise that hinders automated detection. Because the object intensity (the whole cell body) decreases uniformly relative to a static background, comparing Figure. 1 iii) at different defocusing magnitude, a form of negative contrast, akin to shadow fluorescence imaging [17]. One caveat of the Dice score is that adjusting the lateral offset of the image can introduce small overlap errors.

**Figure 1:**
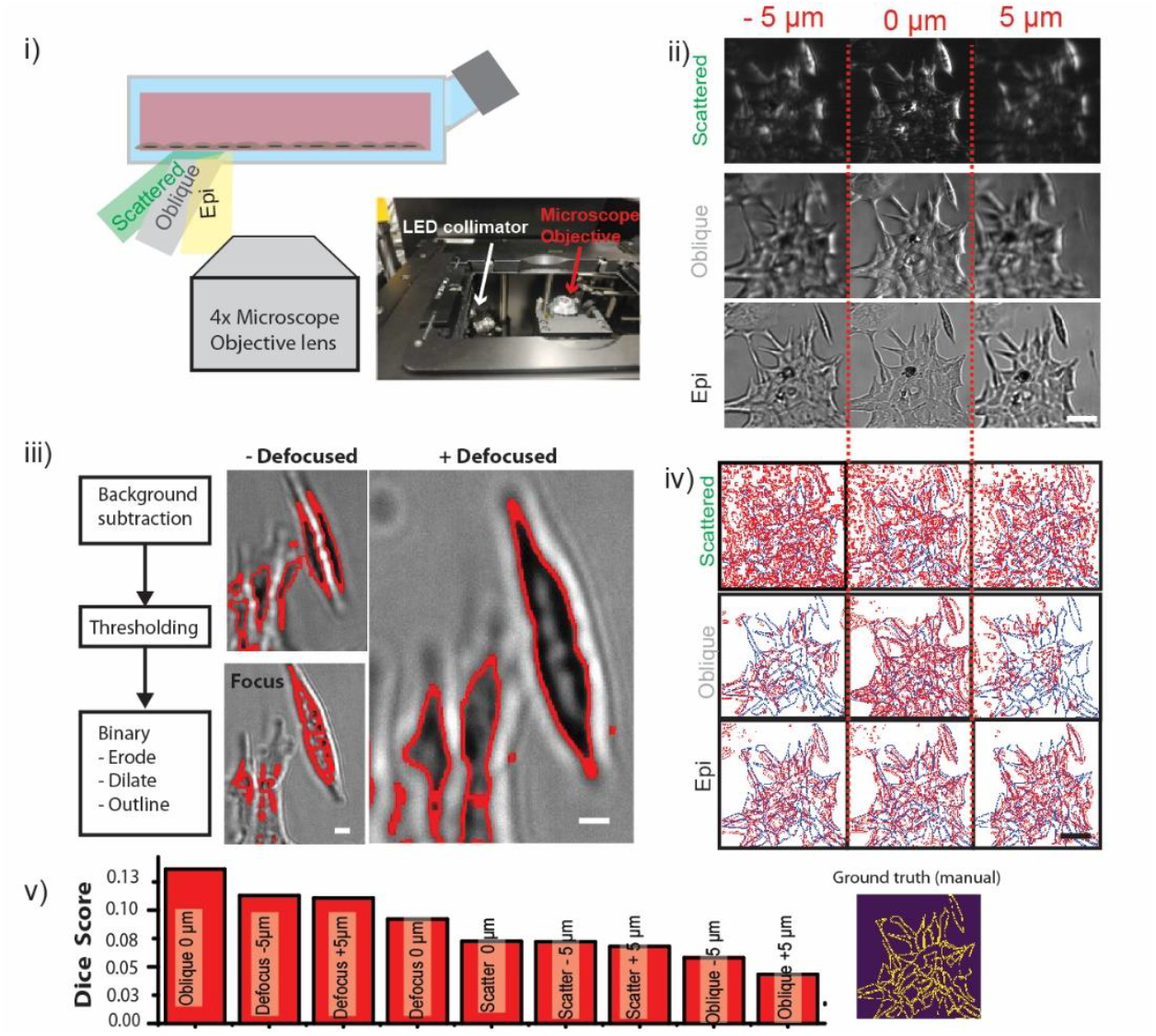
Defocusing Imaging under reflectance illumination. **Comparison of Illumination modes, defocusing images and segmentation performance**. i) Schematic of the optical setup showing three illumination modes: Scattered (green), Oblique (grey), and Epi (yellow), alongside the 4x objective lens with NA of 0.1. ii) Representative images of fibroblast cells and spheroids captured under the three illumination modes at varying focal positions (-5 µm, 0 µm, and +5 µm). iii) Outline the image processing workflow taken (background subtraction, thresholding, and morphological operations (erode, dilate)) in a box diagram alongside with examples of segmented cells and spheroids under epi-illumination. iv) Resulting of segmentation masks (red outline) on the raw image data for each illumination and focusing condition. v) Quantitative ranking of segmentation efficiency utilizing the Dice score. Inset shows ground truth segmented images by manual selection. Scale bar (ii), (iv) 10 µm and (iii) 2 µm

### Does defocus images scale over a large field of view with low numerical aperture optics?

Next, we determine the scalability (field of view) of the defocus imaging. To test this, we require an imaging platform that can operate effective over a large field of view (i.e., millimeter-scale) while still retaining single-cell resolution. In addition to the optical performance (spatial product bandwidth-SBP), we ensured compatibility of the imaging systems that best suited for live cell incubator environments. To do that, we compared two separate platform, an off-shelf digital USB microscope and a home-built microscope system, both placed in separate incubators in shown in <u>Supplementary Figure A1</u> a) i) and ii). Our comparison shown that the digital USB microscope is best suited for large field of view, both in the terms of throughput and form factor for the purpose of live cell imaging. The throughput of a digital USB microscope (based on experimental SBP and number of cells) was shown to be approximately 4-fold higher than the home-built microscope <u>Supplementary Figure A1</u> b) ii). Because the standard off-the-shelf digital USB microscope is equipped with a ring-type reflectance illuminator [18] thus offering uniform illumination under reflectance mode, and can readily fit into a 30 litre incubator.

Since the defocusing mechanical actuators of the digital USB microscopes are crude (+/-20 µm), it is therefore more appropriate to rely on inverted contrast taken from a defocused image instead. As shown in <u>Supplementary Figure A1</u> b) iii), the negative contrast achieved through defocusing with the digital USB microscope reaches a signal-to-noise (SNR) ratio of approximately 1.5 by chose the intensity cross sectional line plot. The SNR ratio of 1.5 was consistent throughout the entire accessible field of view before we set the defocus distance on the USB microscope. Empirically, we observed that the SNR of 1.5 was maintained over several hours to days [19-23] which made it possible to quantify cell number and morphology for each passage using a digital USB microscope.

Next, we proceed to test the use of digital USB microscope in long term cell cultures. Figure 2 shows images of murine cell lines (L929), cancer associated fibroblast (CAFs) and Pancreatic Ductal Adenocacinoma (PDA) grown in flasks (T75 Nunclon Delta flasks) over 16 hours period. Figure. 2a) i) show the full imaging field of view of the digital USB microscope with 80% confluence of murine fibroblast (L-cell-L929, Passage 25). Four individual fields of view (1 to 4) were chosen using ImageJ (ROI manager) and are marked with red and blue circles. Each field of view (FOV) covers an area of approximately ∼ 3 mm^2^. Using ImageJ [24], a macro script was written to count cells by inverting the LUT of the defocused images. The macro encompasses three image processes, as shown in blue rectangular box: inversion of the look-up table (LUT), followed by edge detection, and finally, identification of the maximum intensities to determine the number of cells within each field of view [25]. Fig. 2 a) ii) Display cell counting using four FOVs: raw images (top row), maxima intensity images (bottom row), with counts of 774, 698, 903, and 892 cells. The total number of cell counts over the four selected field of views giving a cell density of 204 cells/mm^2^. A manual check identified an error rate of about 5-8%. A video summary of the macro-output is illustrated in <u>Supplementary Video C1.</u> The reported error in cell counting is less than 5%, which is consistent with other manual and automated cell counting methods [25]. <u>Supplementary Video C2</u> shows approximately 6000 cells imaged on regular cultures T25 flask monitored over 16 hours at 30 second intervals. The aggregated cell count is shown in Figure 2 b) which indicate a growth rate of 1.5 times per day. There were notable noise fluctuations, which can be attributed to changes in illumination intensity. For instance, around the 7.5-hour mark, these variations were likely caused by minor condensation on the LED surface. Longitudinal tracks of cell motility (L929) were conducted on sequence of images from digital USB microscope as shown in Fig. 2 c) using a general cell tracking software known as Trackmate [26] after image segmentation were conducted. Figure. 2 c) ii) and iii) showed two other cell types CAFs and PDA cells imaged at 80 and 20% confluence, video recordings are included in <u>Supplementary Video C3 and C4.</u> Insets ii) and iii) of Figure 2 c) clearly show individual cell bodies based on the inverted LUT map of the defocusing image and after segmentation, thus indicate the suitable to implement defocused imaging on USB digital microscopes that can be broadly used in all existing incubators.

**Figure 2:**
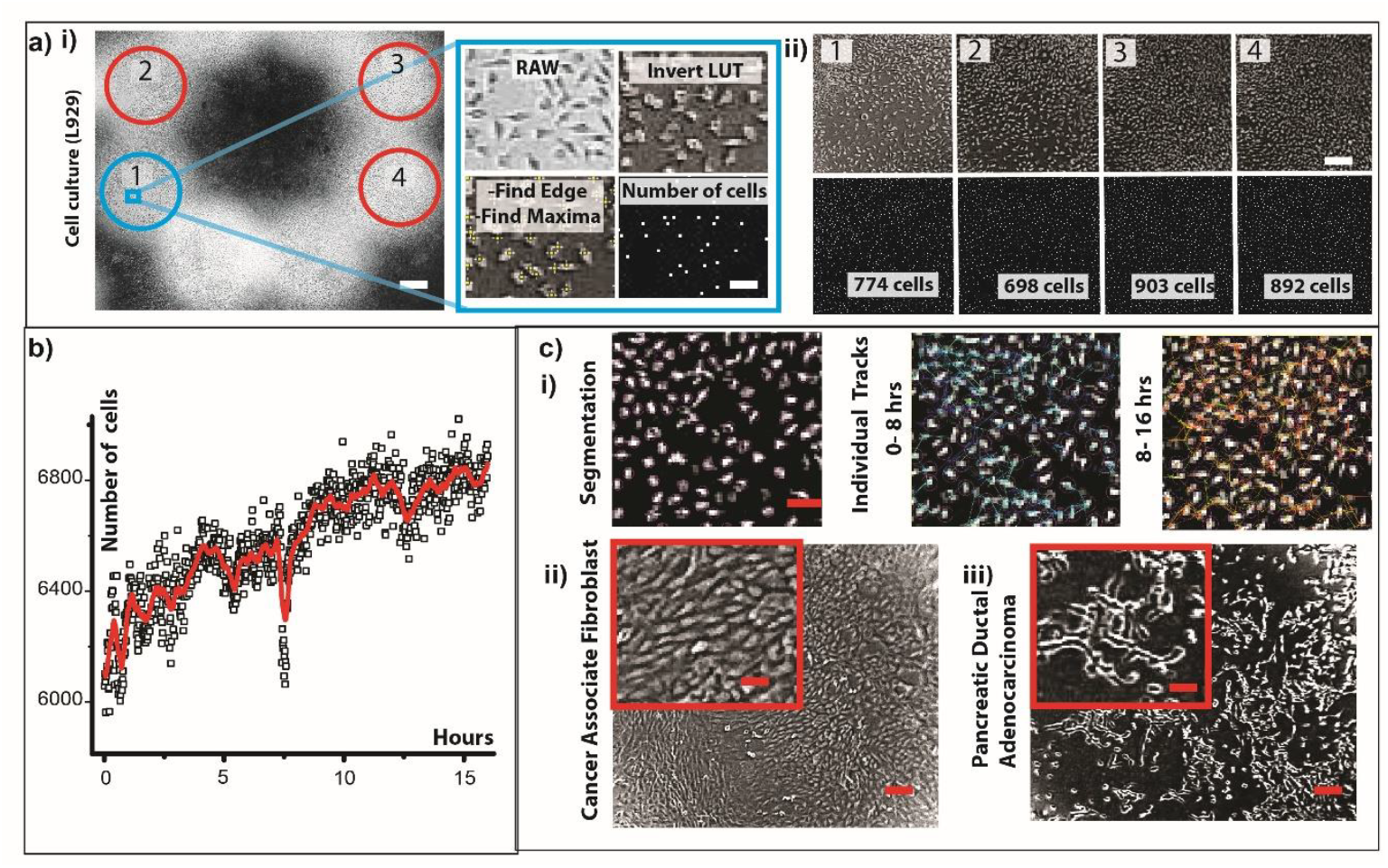
Cell counting and Single cell Tracking using defocus images. a) i) shows the complete imaging field of view captured by the digital USB microscope. Four individual fields of view, circled in red and blue and labeled 1, 2, 3, and 4, were selected for analysis. For cell counting, three post-processing steps were applied: invert LUT, find edge, and find maxima. First, the grayscale map of the raw reflectance image of the cells was inverted. Then, the find edge and find maxima steps were sequentially applied to count the number of cells within each field of view. ii) show the number of cells counted from four region of interest individually at the end of the first 16 hours of culturing. b) shows total number of cells from 1,2,3,4 field of views and the cumulative changes in cell numbers over the same 16 hourr time period. c) show an isolated field of view labelled 1 where images are first segmented, threshold and tracked using Trackmate in ImageJ to determine the individual migration paths of each cell over two 8 hours time interval (0 to 8 hrs and 8 to 16 hrs), c) ii) and iii) shows imaging of highly confluence (70%) cancer associated fibroblast line pancreatic ductal adenocarcinoma cells cells from KPC mice. Scale bar a) i), ii)) 1000 µm, c) i), ii) ii) 200 µm. Inset 50 µm

### Does cell division changes negative contrast derived from defocused images?

Quantitative differences in various adherent cell types such as cell movement and cell division yield functional information about their physical behaviour on different microenvironment. In other words, any changes in cell volume i.e. cell rounding, affects the negative contrast derived from defocus images. We compare the performance of negative contrast taken from defocus images undergoing cell mitosis i.e. cells rounding up. This is important for longitudinal tracking, demonstrating that defocused images from the digital USB microscope possess adequate resolution to quantify longitudinal single-cell activities, including migratory distance and direction as cells undergo mitosis [27]. Figure 3 a) shows two representative images of L929 cells and their tracks after culturing in a T25 flask for a period of 16 hours, observed under a digital USB microscope, Figure 2 a) i) and Figure 2 a) ii) from quantitative phase microscopy [28]. Defocused images of adherent L929 cells taken from digital USB microscope were able to track a total of around 603 cells as shown in <u>Supplementary Video C3</u>. The cell migration tracks recovered from the defocused images taken with the digital USB microscope closely follow the high-quality tracks retrieved from quantitative phase microscopy (minimum tracking distance ∼3 µm), where approximately only 20 cells were tracked. The results show a tracking distance of approximately 172 µm for the digital USB microscope for around 600 cells compared to 160 µm for the quantitative phase microscope taken from 20 cells, indicating comparable tracking performance. At lower confluency, we also examined if defocusing images are affected by different stages of mitosis.

**Figure 3:**
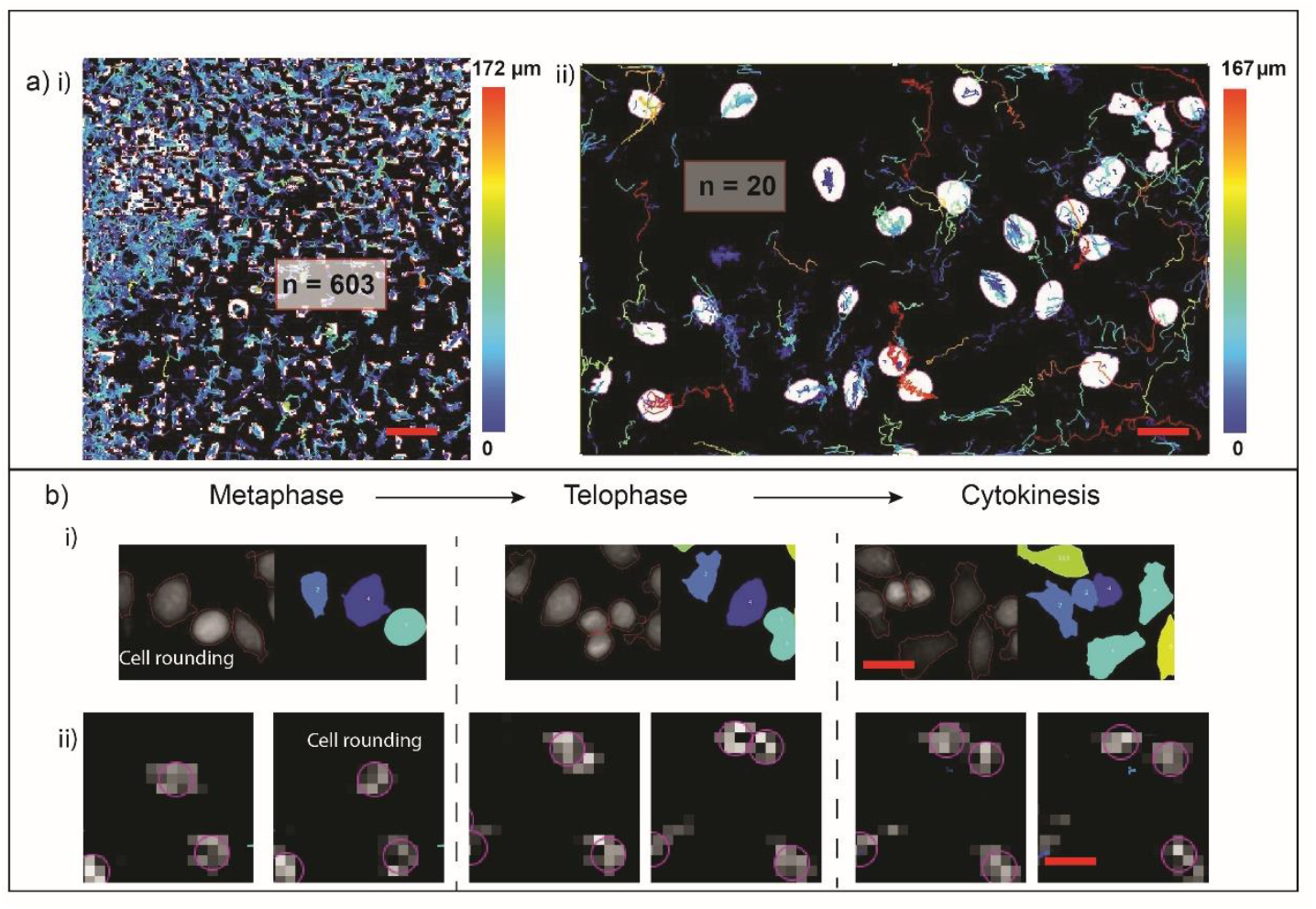
Cell division and proliferation rate) using defocus images. a) Representative images of tracks of individual L929 cell are cultured on T25 Flask and glass bottom dishes i) digital USB microscope - number of cells tracked (n)=603, and 4x objective lens - number of cells tracked (n) =20. Colored tracks indicate displacements. b) i) showing mitosis of cell from a quantitative phase microscope with a 10x objective lens and ii) digital USB microscopy. Scale bar a) i) 100 µm ii)) 50 µm, b) i), ii) 20 µm.

Under high quality quantitative phase imaging system[29, 30], we observed that L929 cells assume distinct morphology of cell rounding [31] at the metaphase stage, as shown Fig. 3b) i). Consequently, under defocused imaging, cell rounding did not affect the negative contrast as originally hypothesis. We can see that negative contrast is maintained during cell splitting/division events can be observed through the temporal tracks as shown in <u>Supplementary Video C5</u> -A video recording of from metaphase to telophase and cytokinesis [32, 33]. This suggests that at a low NA, the long depth of field does not necessarily affect negative contrast captured by action of defocusing; therefore, the negative contrast from defocused images can be used to infer the proliferation rate of the cell population. In the case of L929 cells, we measured a multiplication rate of approximately 1.5–2 times per day, which we validated against other automated cell-counting tools. While we have shown that defocus imaging works for 2D cultures, we have not yet demonstrated its application to 3D cultures [34], which are an increasingly common standard for cell biology. We further examine whether defocused images can serve as an effective image quantification tool for spheroids.

### How does negative contrast of defocus images change during collective spheroid migration and spheroid dissociation rates?

In this section, we investigate the effectiveness of large-field-of-view defocus imaging for tracking 3D tissue spheroid cultures. Since spheroids change its height over time that can alter intended defocus images and so it is necessary to change the imaging lens focal position accordingly. This is also compounded by the fact that light scattering and the physical thickness of these 3D cultures could affect the tracking capabilities derived from negative contrast of defocused images. To evaluate this, we first develop a 3D spheroid culture protocol using L929 cells, as previously shown Fig.1, 2. Our results indicate that L929 cells can express cell adhesion molecules (cadherin) when cultured in Dulbecco’s Modified Eagle Medium (DMEM) without fetal bovine serum (FBS) that subsequently be modulated by adding different MC concentration as <u>Supplementary Figure A2</u>. MC modulates the viscosity of DMEM solution which in turn affects L929 spheroids packing density, shown in <u>Supplementary Figure A2</u>. We called theses modified DMEM as viscous-DMEM (vDMEM). Using this knowledge, we made vDMEM using methyl cellulose 1%, 2% with different viscosities to alter to modulate the rate of spheroid formation and dissociation. Figure 4 a) shows snapshot of defocused images of cell that have been threshold for segmentation at 0 hr before and they aggregated into spheroids after a 40 hr time periods (<u>Supplementary Figure A3, shown 8 hour time interval snapshots)</u>. Figure 4b) i) and ii) show the time series plot of spheroid area and Figure 4 b) i) and ii) circularity appear to increase gradually, indicating consistent growth quadrupling from 200 to 800 µm^2^ and roundness from 0.75 to 0.95. Under all viscosity conditions, spheroid growth, measured by size and circularity, did not vary significantly. Analysis of time lapse imaging also indicates that as the spheroid is increasing its total areas and circularity, each spheroid can independently migrate and the average migration speed of around 12 µm/hr as shown, shown in Fig. 4d) i) and ii). The migration of stromal spheroids might be associated with an increase in cadherin expressions [35]. Figure 4 a) to d) shows shown that defocusing image contrast is maintained for 3D cultured spheroids[36]. Additional details on spheroid preparation sequence in shown in <u>Supplementary Figure.A4.</u>

**Figure 4:**
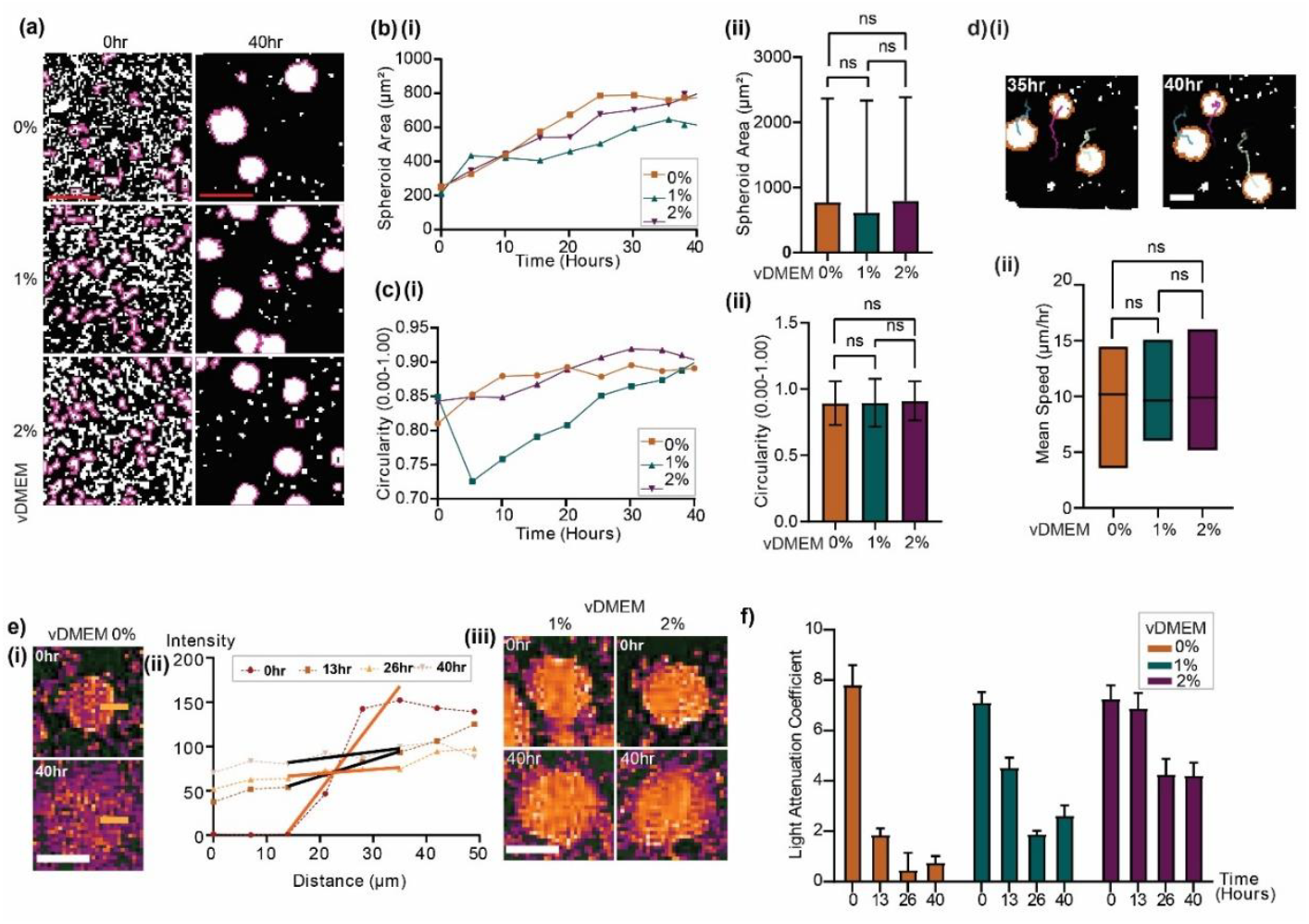
Quantifying changes to defocusing contrast with 3D spheroids. **(a) i)** Representative images of the use of circularity filter from TrackMate Filter (Circularity ≥0.23-criterion) used to extract spheroids from cell cultured in all vDMEM conditions at 0hr and 40hr. Each purple outline identifies cell aggregates which met the criteria. **(b)(i) and (ii)** Longitudinal changes to the size of aggregate (μm^2^) and circularity across the 40-hour culture for vDMEM conditions (N spheroids=12). Each data point are calculated mean values from 12 spheroids. **b)ii)** Final size of aggregate (μm^2^) after 40-hours for 0% (n=431), 1% (n=575) and 2% (n=569) vDMEM. Unpaired T-tests indicated no significant difference in final aggregate size between vDMEM conditions. Data represented as mean±s.d. **(c)(i)** Circularity (0.00-1.00) of individual spheroids across 40-hour culture for all vDMEM conditions (N spheroids=12). Data represented as mean values of 12 spheroids. **c) (ii)** plots the final circularity (0.00-1.00) of aggregates after 40-hour culture for 0% (n=656), 1% (n=714) and 2% (n=696) vDMEM. Unpaired T tests indicated no significant difference in final aggregate circularity between vDMEM conditions. Data represented as mean values±s.d. **d)i)** Representative images of spheroid migration over ≥10 hours for 0% DMEM at 35hr and 40hr. Orange outline identifies spheroids and corresponding migration paths (purple, green lines). **d) ii)** Average speed of spheroids (μm/hr) between 30 - 40hrs for vDMEM conditions 0% (n=42), 1% (n=29), and 2% (n=35). Unpaired T-tests revealed no significant difference in mean speed of spheroids travelling under all the vDMEM conditions. **e) i**) Dissociation of stromal spheroids is marked by the intensity gradient (inverted LUT) measured along the orange line (50μm in length). **e) ii)** Plot of raw intensity along with a linear regression (thick line – black and orange). Simple linear regression was used to fit intensity gradient (distance 14-35μm) (e.g. Slope = 7.178, R2 = 0.9745). **e) iii)** shows intensity of spheroids developed in all the vDMEM immediately (0hr) and 40-hours (40hr) after resuspension resuspended in DMEM+FBS. **f)** Light attenuation coefficient (inverted LUT) from linear regression resuspension for spheroids previous developed in vDMEM across 0hr, 13hr, 26hr and 40hrs. All profiles had R-squared value of ≥0.8, except from 0% at 26hr and 0% at 40hr. Two-tailed p-value of slopes indicated that difference of slopes within vDMEM conditions 0% (p<0.0001), 1% (p<0.0001) and 2% (p=0.0007) were statistically significant. Data represented are mean±s.d. a) Scale bars represent 200μm. d), e) 100μm.

Next, we identified if contrast of defocusing imaging could change when each of the spheroid begin to dissociate, which is useful for anticancer drugs or immune infiltration studies [37]. The reason is that as spheroid dissociate, there is likely more optical challenges particularly optical turbidity i.e. uneven contrast changes over time. To measure changes in contrast, we extracted the spheroids and resuspended them into DMEM that contains FBS. As shown in Figure 4e i) and ii) over a 40 hr period during dissociation, an intensity gradient was used to determine the changes in image contrast <u>(Supplementary Figure A5)</u>. The intensity is extracted from the yellow line plot of a chosen spheroid versus the background as shown in Figure 4e) i) and Figure 4e ii) shows the changing negative contrast follows a linear intensity attenuation coefficient from this plot. Contrast decreases with the spheroid dissociation. Lower coefficients indicate a faster reduction in imaging contrast, primarily signaling the complete dissociation of the spheroids. Figure 4e) iii) shows two other spheroid formed at vDMEM 1% and 2% that are resuspended into DMEM and FBS solutions over 40 hours period. Figure 4 f) summaries the spheroids and the changes to the contrast based on the light attenuations coefficient. Over the course of 40 hours, we observed the average contrast of the defocused spheroid images does reduces the rate of dissociation rate of the 1% and 2%, thus indicate a change in thickness. The change in negative contrast indicates an increase in biological cell-cell adhesion at the spheroid core. [17, 38, 39].

## Conclusion

Our results showed that defocusing imaging can be a highly valuable tool in live cell imaging that offers sufficient imaging contrast and resolution across their field of view. This approach is scalable, as a rudimentary digital USB microscope can readily utilize defocused images for routine cell culture, spheroid formation, and cell invasion assays [40, 41] without resorting to advanced 3D imaging tools. Unlike computational imaging technologies such as lensless microscopy [42] and Fourier Ptychography [43], defocusing imaging does not require complex computational approaches and is immediately compatible with thick imaging polymer walls of standard cell culturing flasks. The defocusing method is suitable for a range of systems, from compound microscopes to simple, digital USB devices equipped with all necessary illumination and imaging sensors. Furthermore, the software drivers (OpenCV) can be directly integrated with the open-source Micro-Manager program for single-cell imaging. Our results further suggest that the defocus images of cells and spheroids captured by a digital USB microscope can be digitally processed to investigate single-cell and spheroid dynamics, from cell migration and division to spheroid formation and dissociation. Next, we aim to expand the use of multiple digital USB microscopes for capturing multi-well tissue culture plates, this will permit parallel imaging simultaneously. Commercial or open-source *incuscope* system [44] could leverage the importance of defocusing imaging using lower numerical aperture optics to expand their applications into spheroid imaging. Additionally, in <u>Supplementary Figure A6</u> and <u>Supplementary Table B1,</u> we provide the comparison of different incubator-compatible microscope technologies in terms of throughput, price and spatial product bandwidth for subcellular to single cells and whole organoid.

## Acknowledgement

We would like to thank Melanie White from University of Queensland for the critical comments on the manuscript. W M Lee acknowledges Hari Shroff from Janelia Research Campus for pointing out to us the history of defocusing microscopy.

## Methods

### Protocol to prepare Methyl-Cellulose (MC)

Viscous DMEM was prepared through addition of 41kDa Methylcellulose (Sigma Alrich) into 1X DMEM (Low Glucose Dulbecco’s Modified Eagle Serum 1X, Gibco). Media was prepared in batches of 75mL, at 1% and 2% concentration of Methylcellulose (MC). To prepare media, 1/3 of the total volume of 1X DMEM was transferred to a 50mL tube and prewarmed to 80°C for 30-minutes. The remaining 2/3 of DMEM was stored in fridge at 4°C. MC was weighed to desired % w/v for total media. Prewarmed DMEM and MC were transferred to 100mL bottle and placed on 80°C magnetic stirring plate for 30-minutes. Separately, a 5L beaker was prepared with 500mL of 4°C tap water and supplemented with a suitable amount of ice to maintain it at 4°C. After 30-minutes, the hot plate was turned off and the remaining 2/3 of DMEM media was transferred to the vDMEM mixture. The vDMEM bottle was immediately immersed in the 5L beaker and returned to the magnetic stirrer for 3 hours. The temperature of water in the 5L beaker was checked at 30-minute intervals using a mercury thermometer to ensure it did not deviate from 4°C. After 90 minutes the water was supplemented with additional ice/ freezer bricks. Once complete, the vDMEM media was transferred to fridge at 4°C until filtration. Centrifugal filtration was performed to separate large molecules of undissolved MC using gravitational force. vDMEM media at 4°C was aliquoted into 6 1.5mL Eppendorf tubes using a 5mL serological pipette and centrifuged at 21300 x g for 50minutes. 1.5mL Eppendorf tubes were used as designed for our centrifuge and rotor capacity. The supernatant from each Eppendorf tube (approx. 1.3mL of media) was removed using a different 5mL serological pipette and transferred to a 50mL tube. The process was repeated until all the unfiltered vDMEM media had been centrifuged. The same set of centrifuge tubes were used for the entire bottle. A new 50mL tube was acquired for the supernatant once filled. Centrifugal filtration was followed by an additional mechanical filtration step in the laminar flow cabinet. The 50mL tubes containing centrifuge filtered MC-Media were sterilised and placed in cabinet, as well as 2x 10mL syringes, 2x 0.45-micron syringe filters, 2x 0.2-micron syringe filters, and 2x empty 50mL tubes. A 0.45-micron filter was attached to a 10mL syringe and placed on top of an empty 50mL tube with the lid removed. After removing the syringe plunger, approximately 6mL of vDMEM media was transferred into the syringe barrel. The syringe plunger was then added back to the barrel to seal, and the syringe with filter attached was carefully inverted. With the syringe inverted, the filter was loosened and plunger pushed slowly to remove air bubbles in the syringe cavity, being careful to avoid contact with any extruded media. The filter was then tightened, and syringe placed upright over the 50mL tube and media filtered through. Removing the air bubbles reduces air pressure resistance through the filter. This process was repeated until all media was filtered through the 0.45-micron filter. The filter and syringe were replaced every 2-3 repeats as filters became blocked. The 0.45-micron filtered vDMEM media underwent additional filtering through a 0.2-micron filter into a new 50mL tube following the same filtering procedure. Mechanical filtration served as an observational confirmation of increased viscosity, with increased resistance through the filter indicative of successful thermosensitive setting of vDMEM. Furthermore, the mechanical filtration removes small particles of debris or undissolved MC that may be present in the vDMEM.

### Protocol for spheroid formation

L929 cells for spheroid formation were harvested from the primary passage flask during cell passaging and cell count taken note. Detached cells were centrifuged for 5-minutes at 300 x g in a 15mL tube, and the supernatant (DMEM+FBS) was discarded. Cells were resuspended in the serum-free vDMEM with the volume determined by the original cell count, for consistent seeding at 40% confluency (∼2.5x10^6^ cells) in a 25cm^2^ flask (Nunc EasYFlask # 156367, ThermoFisher). Cells were seeded in 7mL of the prepared the viscous DMEM media or DMEM only (for the viscosity control) at 37°C with 5% CO_2_. Spheroid formation was monitored over 40-hours using live cell imaging

### Protocol for spheroid dissociation

To study spheroid dissociation, serum free vDMEM media was removed from the flask and refreshed with 7mL DMEM+10% FBS. All spheroid containing media was removed in 1mL increments using micropipette and transferred to a 15mL centrifuge tube. 1mL volume transfer was necessary to reduce loss of spheroids to the pipette. The spheroid containing media was centrifuged for 10-minutes at 600 x g and the vDMEM supernatant was removed. Spheroids were resuspended in 3mL of PBS and centrifuged again for 5-minutes at 300 x g to wash, repeating this process of removal and resuspension in PBS 3 times. The original T25 flask was washed with 3mL of PBS 3 times. After washing, 6mL of prewarmed DMEM+10% FBS media was transferred to the original T25. Spheroids were resuspended in 1mL of DMEM+ 10% FBS, using mechanical aspiration to redisperse throughout media. Spheroids were transferred back to the T25 flask using a 500µL micropipette. Flasks were checked under standard brightfield to ensure spheroids were intact. Spheroids were monitored over 40-hours using live cell imaging in standard incubation of 37°C with 5% CO_2_.

## Supplementary Files

A. **Supplementary Figures**
  - **Figure A1 - Determining suitable compact microscope for optimal throughput to study defocus imaging in live cells**
  - **Figure. A1 - Cadherin expressed in L929 spheroid after culturing in 0%, 1%, 2% vDMEM without fetal bovine serum**
  - **Figure A2 - Representative images of spheroid formation with vDMEM 1%, 2%**
  - **Figure A3 - Intensity gradient (light attenuation coefficient) measured for each vDMEM solution**
  - **Figure A4 – Spheroid formation and dissociation experiment**
  - **Figure A5 - Intensity gradient (light attenuation coefficient) measured for each vDMEM solution**
B. **Table B1- – Comparison of incubator microscope**
C. **Video C1 – Cell Counting** **Video C2 – Imaging L929 cells over 16 hours at 60 second intervals** **Video C3 – Single cell tracking L929 migration over 3hrs using trackmate Video C4 – Seeding KPC cells on T75 flask** **Video C5- Mitosis of L929 with quantitive microscopy and digital USB microscope** **Video C6 – Tracking of spheroid movement in 1% vDMEM**

