## supplementary figures and table for "Defocused reflectance imaging for low numerical aperture, large field of view quantitative live cell imaging studies"

**A) Supplementary Figures**
- **Figure A1 - Optimal throughput and defocusing contrast**
- **Figure. A1 - Cadherin expressed in L929 spheroid after culturing in 0%, 1%, 2% vDMEM without fetal bovine serum**
- **Figure A2 - Representative images of spheroid formation with vDMEM 1%, 2%**
- **Figure A3 - Intensity gradient (light attenuation coefficient) measured for each vDMEM solution**
- **Figure A4 – Spheroid formation and dissociation experiment**
- **Figure A5 - Intensity gradient (light attenuation coefficient) measured for each vDMEM solution**

**B) Table B1- – Comparison of incubator microscope**

**C) Supplementary Videos**

**Video C1 – Cell Counting**
**Video C2 – Imaging L929 cells over 16 hours at 60 second intervals**
**Video C3 – Single cell tracking L929 migration over 3hrs using trackmate**
**Video C4 – Seeding KPC cells on T75 flask**
**Video C5- Mitosis of L929 with quantitive microscopy and handheld microscope**
**Video C6 – Tracking of spheroid movement in 1% vDMEM**

**Figure A1 - Optimal throughput and defocusing contrast**

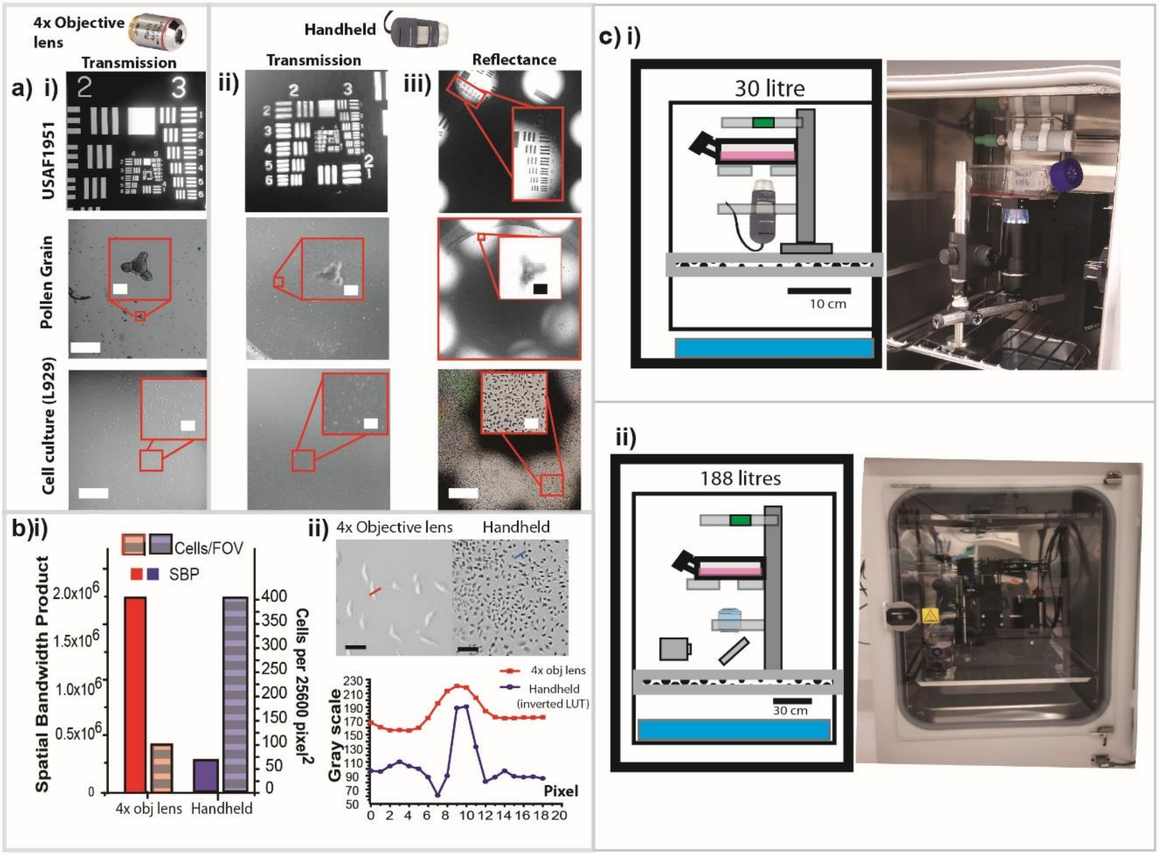


Comparison of images quality, spatial bandwidth product and throughput of commercial microscope objective lens (Nikon TE2000, 4x objective lens and 10x eye piece magnification, 8 megapixels-OV08B) and handheld microscope (Digitech, 5MP USB Digital Microscope) with fixed and live samples- USAF1951, fixed pollen grain and live stromal cells (L929) cultured in T75 cell culture flask. The home-built microscope in transmission mode was used for comparison, utilizing the same 4x objective lens with a numerical aperture (NA) of 0.1 (Fig.A1) without any tube lens (making it non-infinity corrected). Fig. A1 show the imaging performance of two types of microscopes, where we determined optimal throughput by the total number of cells that are resolved in a single frame. For an efficient live cell imaging system, one should allocate the minimum number of pixels over the entire field of view, also known in optical system as a dimensionless unit called spatial product bandwidth, SBP [1].

Figure A1a) shows each imaging system conducted with a USAF 1951 card, fixed pollen grain and live adherent cells in a thick culture flask for both microscope system (4x home built and handheld microscope), which was used to calibrate the accessible field of view and experimental SBP. Figure A1a) i) shows the imaging resolution of the 4x objective lens is able to provide a SBP of 2.7x $10^6$ based on a single resolvable spot of 3.36 µm with a field of view of 5.17 mm (full field of view not shown- due to size limitation). In comparison, Fig. A1a) ii) the transmitted imaging images from handheld microscope was able to cover a total field of view of 7.619 mm with resolvable features of 8.77 µm, approximately NA of 0.033. measures at 3.91 µm with a 4x objective lens. On the other hand, for this specific handheld microscope model from Digitech shown in Figure. A1a) iii), under reflectance mode, the field of view was reduced to 5.750 mm whilst maintaining a lateral resolution of 8.77 µm. As expected, the SBP of transmission brightfield microscope is 3 folds higher than handheld microscope in Figure. A1 b) i). Nevertheless, both imaging system can clearly resolve a single pollen grain and single adherent cells. Comparing defocusing imaging from home-built microscope and handheld microscope, the low magnification handheld microscope utilises only 64 $pixel^2$ (8 x 8) which is 6 folds less pixels than the 400 $pixel^2$ used in a transmission brightfield microscope. Figure A1 b) ii) and iii) shows that for the same pixel area of around 25,600 $pixel^2$, the throughput of a handheld microscope is already 4-fold higher than the microscope objective that is optimal. Figure A1b(ii)(top) displays the number of adherent cells captured within a 25,600 $pixel^2$ area using both a 4x microscope and a handheld microscope. Figure A1b(ii)(bottom) presents a cross-sectional intensity plots the defocusing imaging contrast of handheld microscope (signal to noise ratio- SNR ~1.4) against the defocused transmitted imaging contrast of 4x objective lenses (SNR ~0.4). The line plot of the reflectance image intensity of the handheld microscope is inverted for comparison. Because of the small form factor, the handheld microscope can easily fill into a compact incubator of only 30 litres. In comparison, the microscope using 4x objective requires an incubator of a size of 188 litres to fit, as shown in Figure. A1c) i) because of addition of condenser and tube lens. Scale bar a) i), ii), iii) _1000 µm, inset 50 µm. b) ii) 50 µm. Since our study focuses the immediate application in cell biology, we did not address optical factors that influence imaging throughput, such as intermediate magnification lenses and the pixel size of the camera sensor.

**Figure. A2 - Cadherin expressed in L929 spheroid after culturing in 0%, 1%, 2% vDMEM without fetal bovine serum**

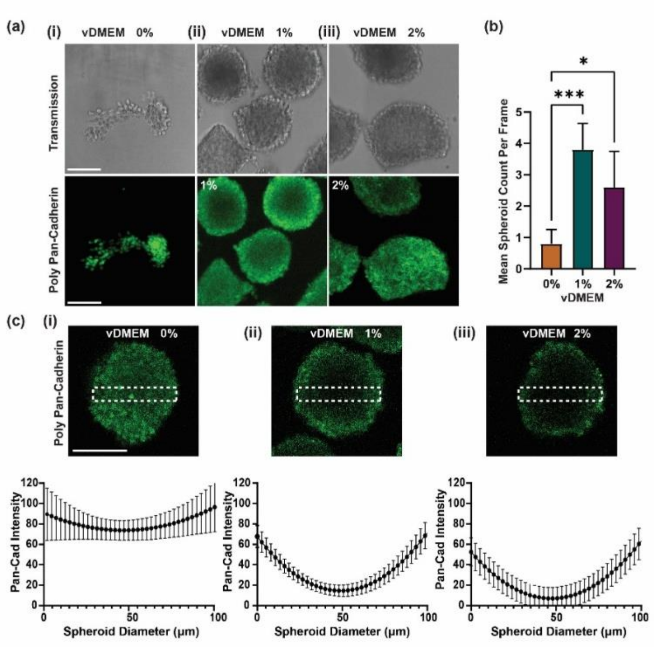


**Figure A2.** Immunocytochemistry staining *(Pan-cadherin Primary antibody - Invitrogen, Cat # 71-7100, Pan-cadherin Polyclonal Antibody, Secondary antibody - Abcam, Ab150065, Donkey anti-Rabbit IgG H&L Alexa Fluor 488 preabsorbed)* were used to measure the cadherin expression of L929 spheroids. After 4-day culture, spheroid samples were centrifuged for 5-minutes at 300 x g for 20 times prior to staining and labelling. Transmission and fluorescence images of L929 spheroids are shown in **Figure. A1 a)i), ii), iii).** Fluorescence images confirmed that L929 spheroids readily express cadherin when cultured in DMEM alone and without FBS. Maximum projection of pan-cadherin expression over 60μm depth. **Figure. A1) b)** Determine the number of spheroid expresses Pan-cadherin. Mean number of spheroids detected per frame during confocal imaging. Spheroids were developed over 4 days before fixing. Data represented as mean±s.d. Unpaired *t*-tests determined a statistically significant difference in the mean counts of 0% and both elevated viscosity conditions 1% (p=0.001) and 2% (p=0.0111). **(c)** Expression of pan-cadherin at cross-section of spheroid centre for varying vDMEM conditions. Spheroids of consistent diameter were used (~160μm). The rectangle represents region used for intensity plot profile, with the length equal to the diameter and height 1/4 of the diameter. Scale bar represents 100μm. **(b)** Mean Pan-cadherin fluorescent intensity distribution across spheroid diameter (normalised to 100μm) for varying vDMEM conditions (n=3). Using Ordinary One-Way Anova the difference between intensity profiles was deemed extremely significant (p<0.0001). Data represented as mean±s.d **c)** Maximum Z-Projection intensity over 60μm indicated difference of fluorescence distribution show different intensity profiles across the spheroids, suggesting pan-Cadherin expression that indicate higher packing density for spheroids in 1% and 2% vDMEM
a), c) Scale bars represent 100μm.

**Figure. A3 - Representative images of spheroid formation with vDMEM 1%, 2% over 40 hours**

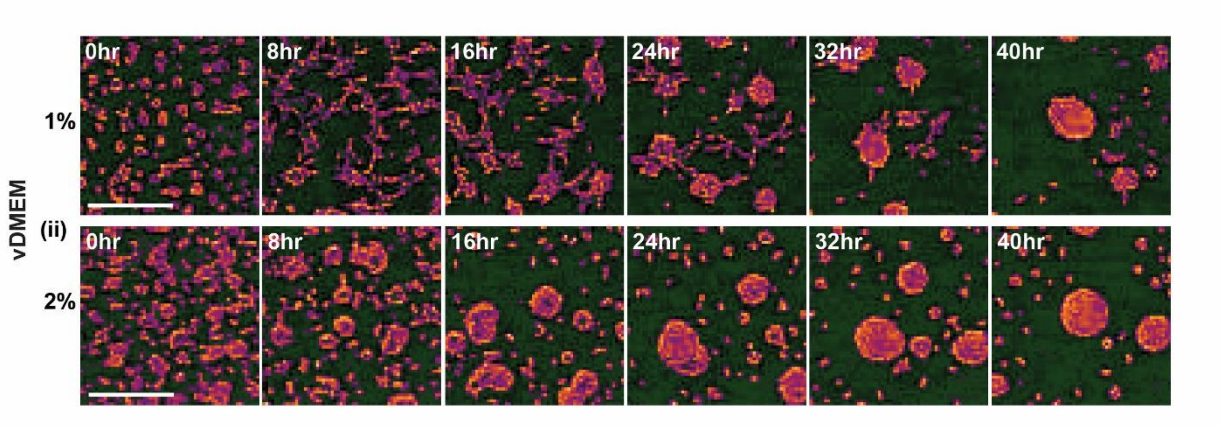


**Figure A2**_Morphology of aggregates across 40 hours for **(i)** 1% DMEM and **(ii)** 2% DMEM in 8-hour intervals. Scale bar represents 200µm.

**Figure A4 – Spheroid formation and dissociation experiment**

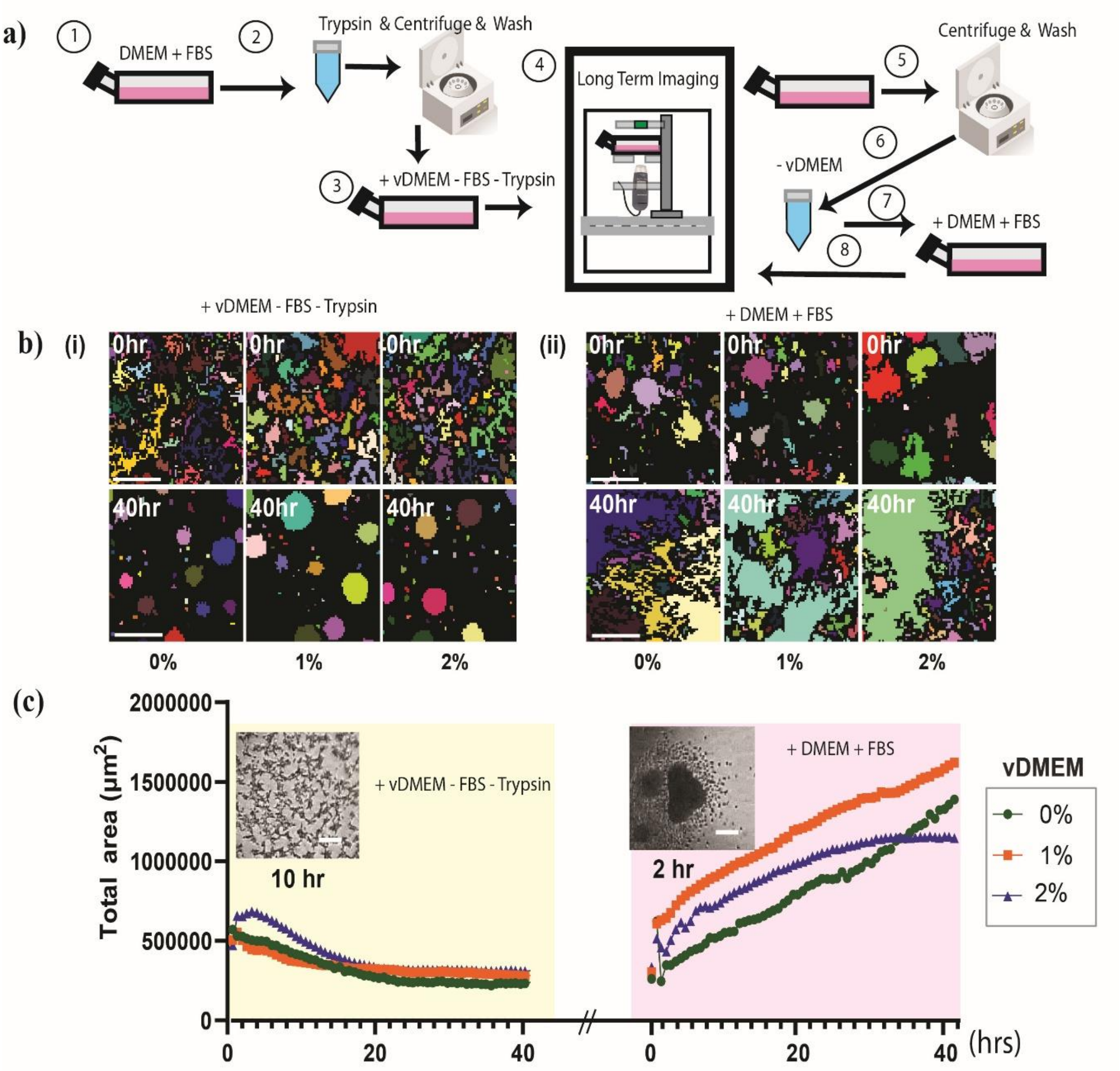


Illustrates the sequence of spheroid preparation. L929 cells were initially harvested from standard culture media (DMEM with 5% FBS) once they reached full confluence. They were then subjected to trypsinization, centrifugation, and washing before being re-seeded into viscosity DMEM (v_DMEM) without FBS. Two vDMEM solutions were made using MC concentrations of 1%, 2% respectively (Supplementary Protocol). After extended imaging and spheroid formation, the vDMEM culture is centrifuged and washed to remove vDMEM. The culture is then resuspended in regular culture medium (DMEM + 10% FBS) for long-term monitoring of spheroid dissociation. Figure. 4 b) i) shows images taken from handheld microscope at Step 4 under different vDMEM solutions. The initial number of seeded cells is approximately 781,250 cells for a T25 flask. Figure 4b) i) also shows L929 cell readily organise into spheroids from 24 to 41 hours (1.7 days). Initial surface area measurements of the total number of cells show that cells from vDMEM-0, 1%, and

2% all aggregate into distinct spheroids within the first 24 hours. Figure 4b) ii) shows that cells in resuspended spheroids with DMEM+FBS began to spread and occupy space in the culture flask. Figure 4 c shows the surface area measurement in a single plot where the cell-cell aggregation occurs in the first 20 hours without FBS in all three different vDMEM solutions, whilst within the first 2 hours of being resuspended in DMEM+ FBS, the spheroid begun to dissociate rapidly. The dissociation rate indicates the difference of spheroid behaviour cultured in the different vDMEM solutions. We observed that with 0 to 1% vDMEM, the disassociation was similar, but at 2% we observed the total area reach a plateau. It appears that 3D spheroids can be readily imaged using the same image processing steps. b) representative image of the initial (0 hrs) and final conditions (40 hrs). Color indicate segmentation. c) time series plot of the total area quantified over 41 hours.

Scale bar b) i), ii)) 100 µm, c) inset 100 µm

**Supplementary Fig. A5 - Intensity gradient (light attenuation coefficient) measured for each vDMEM solution**

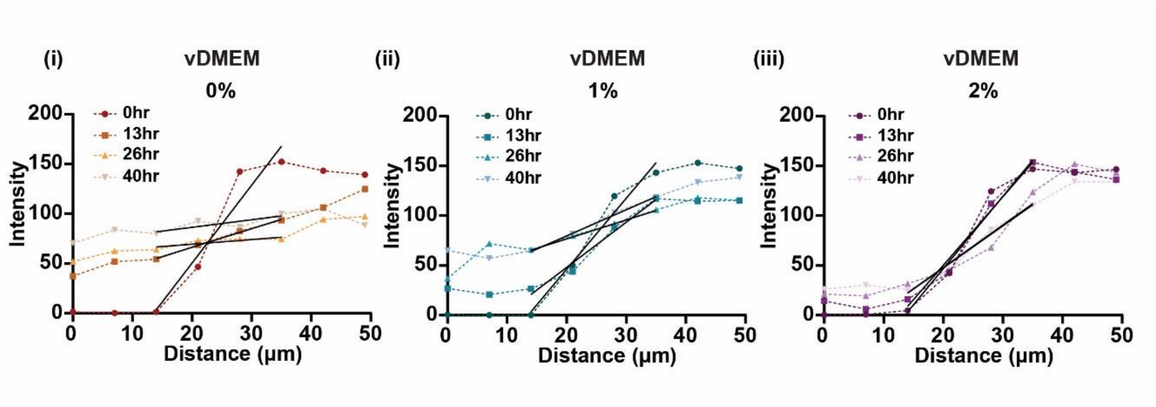


**Intensity gradient (light attenuation coefficient) used to measure changes in negative contrast from defocus images for** dissociating spheroids under different vDMEM conditions. 500 image stacks of 1280x1024 pixels (8963.59 x 7170.87 µm) FOV were reduced to 4 to reflect the 4 times points of spheroid dissociation. The images were then normalized to resolve differences in exposure. Manual detection of spheroids of near equal diameter (90-110µm) was used for intensity profiles as an accurate comparison between vDMEM conditions. A 50 µm plot profile was taken across the border of the spheroid, with the spheroid-background border identified in the first frame and maintained at the ~25µm mark across all time frames. This required manual adjustment to account for movement of spheroids within the frame. Plot profile values were processed using GraphPad Prism.

**Supplementary Figure A6 – Comparative Advantages of cellular imaging devices**

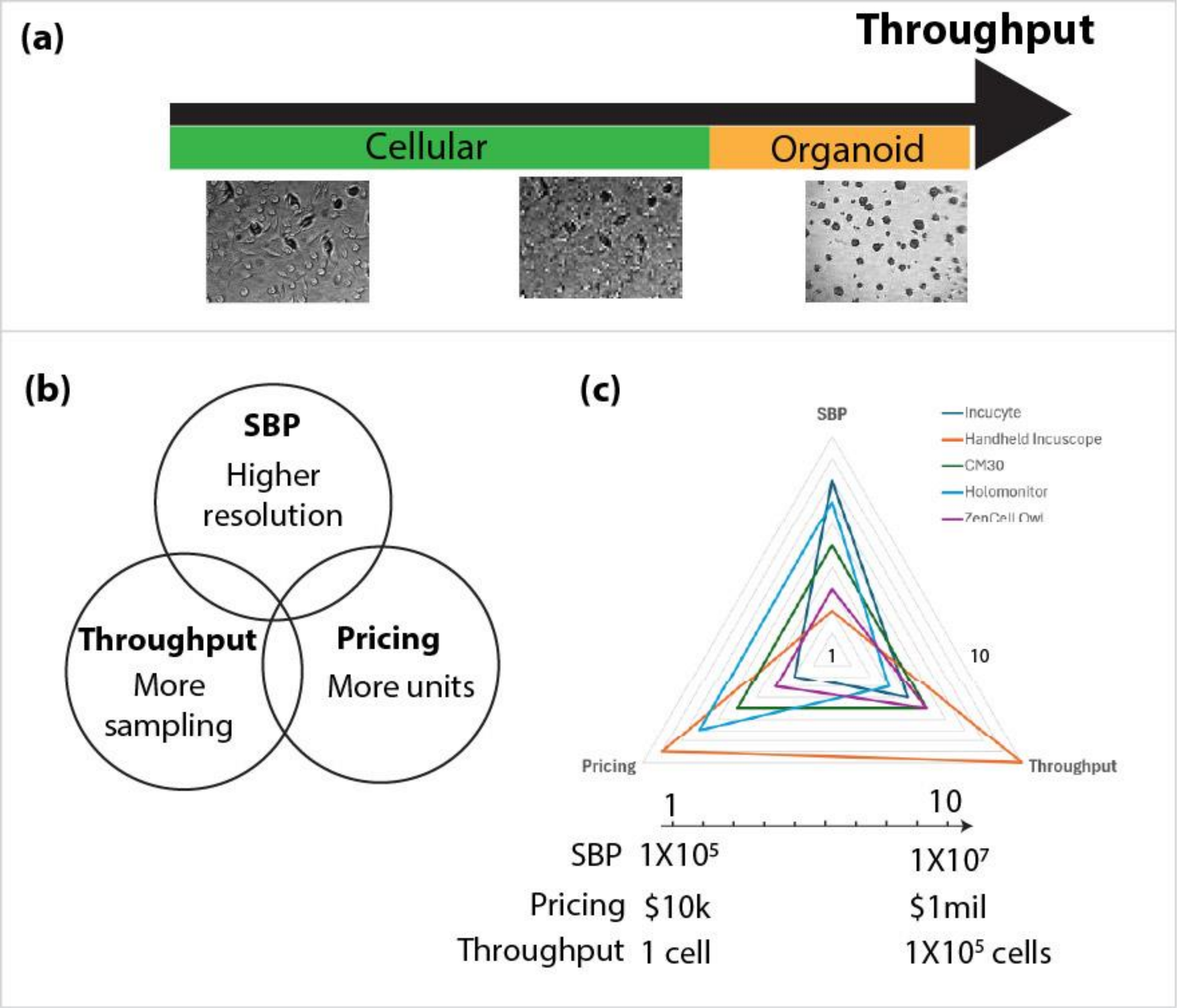


Qualitative evaluating performance of cellular to organoid imaging through SBP, sample throughput and pricing of devices. a) Individual cells to organoids are distinct in images when represented by sufficient number of pixels. Lowering this number increases throughput at the cost of resolution. b) Decision points for balancing between SBP, throughput and pricing of incuscopes for cell imaging. c) Radar graph comparing SBP, throughput and pricing of different commercial incuscopes in market and handheld incuscopes used in this report.

**B)**

**Table B1 – Comparison of incubator microscope**

| Microscope | Cost | Integrated lighting | Industry package | microManager Compaitbiliy |
|---|---|---|---|---|
| DIGITECH (paper) | AUD$300 | Yes | Yes | Yes |
| INCUYTE | AUD$25,000* | Yes | Yes | No |
| CM30 | AUD$10,000* | Yes | Yes | No |

*- estimates

**C1 Video 1 – Cell Counting**

Screen video recording of cell counting using handheld microscope with micromanager and macro – maxima_macro_cellcount using inverted LUT of defocus images

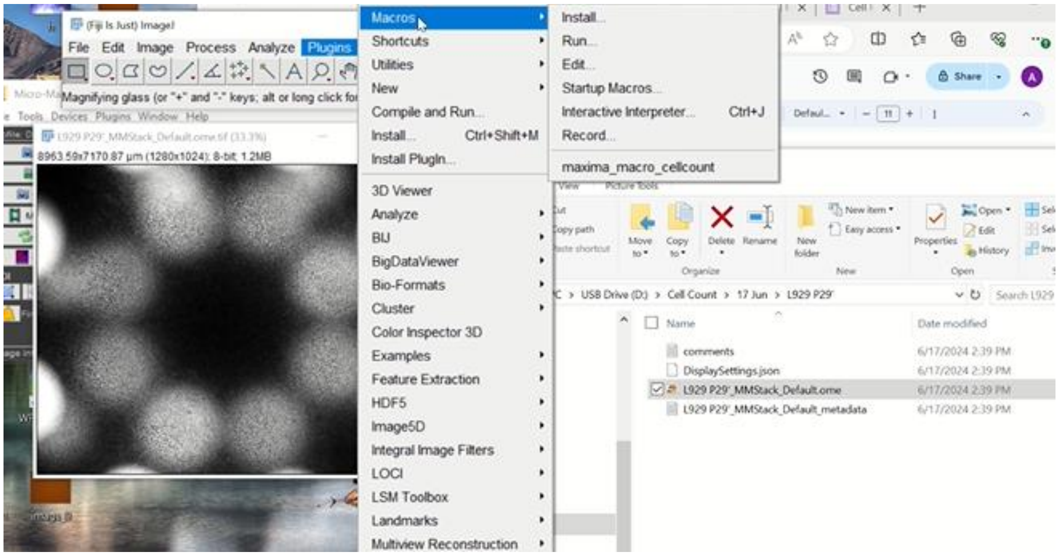

**Video C2 – Inverted LUT of defocus L929 cells over 16 hours at 60 second intervals**

Animated sequence of images of L929 cultured in DMEM+FBS in T75 Flask over 16 hours at 1 minute time intervals

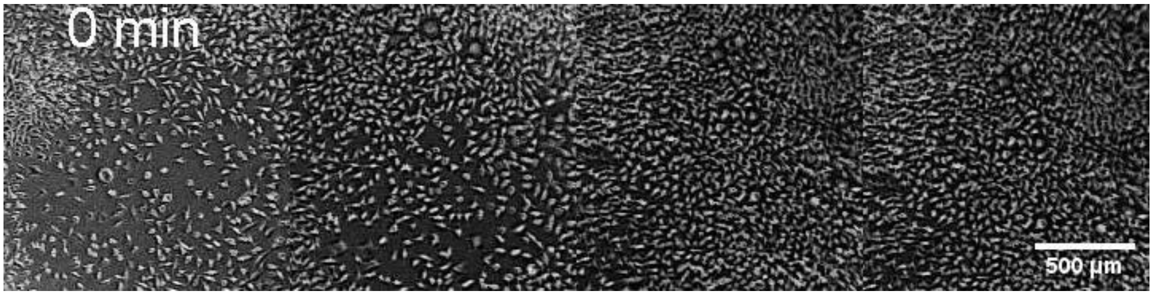

**Video C3 – Single cell tracking L929 migration over 3hrs using trackmate**

Animated sequence of images of tracking L929 migrating in T75 flask over 3 hour time period

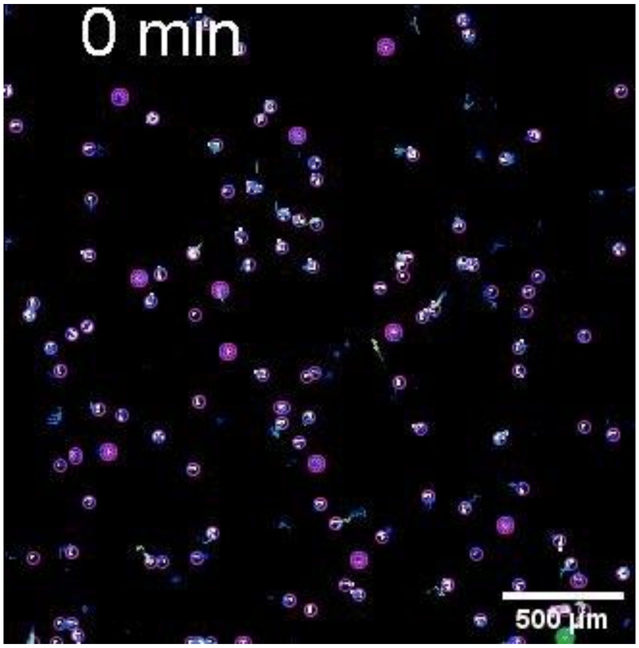

**Video C4 – Seeding KPC cells on T75 flask**

Animated sequence of images of KPC cells seeded in T75 flask forming long-chained tissue like networks (false color code- green on inverted LUT)

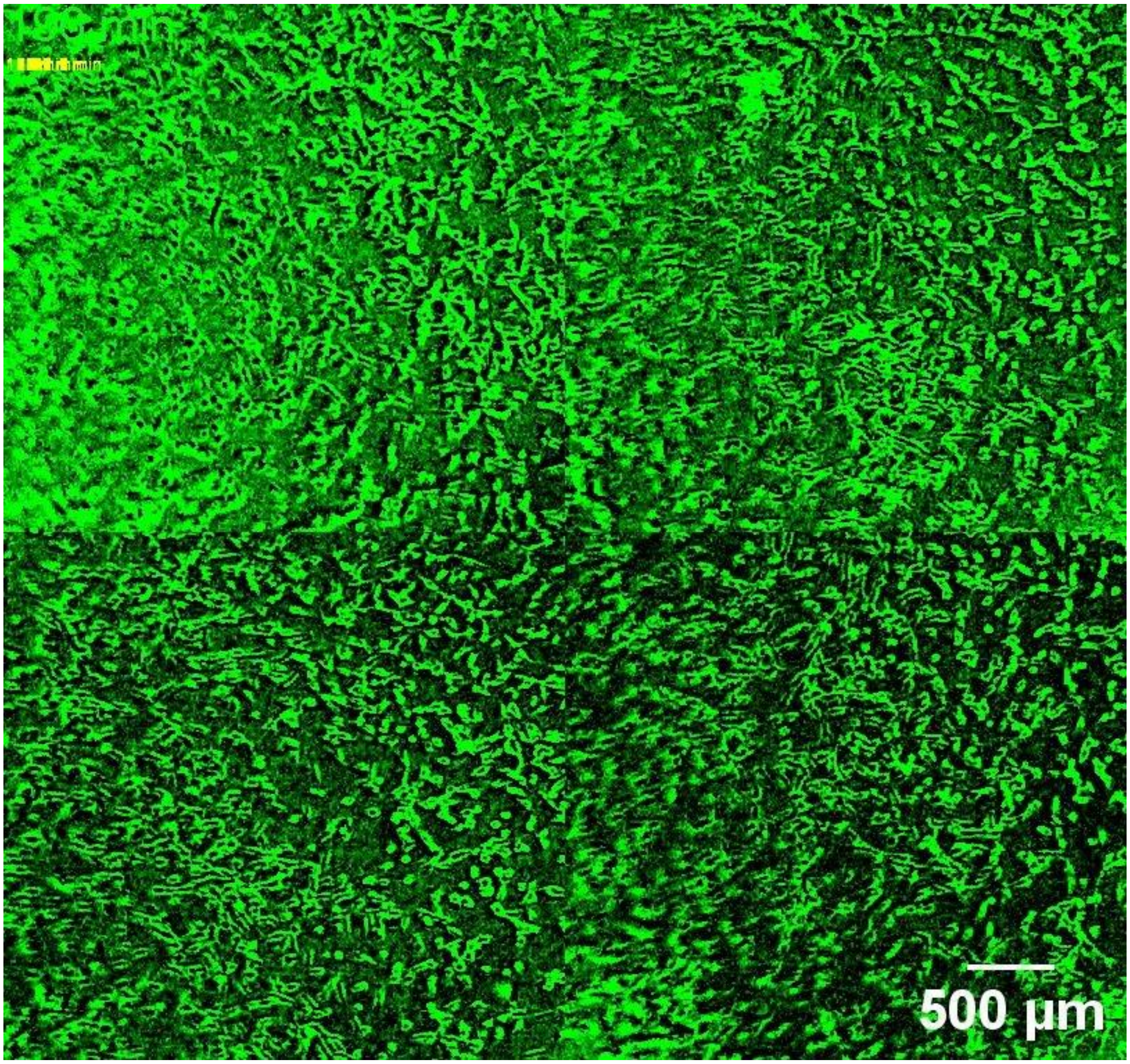

**Video C5- Cell Divison L929. Comparing quantitive phase microscopy (QPM on the left) and inverted LUT (handheld microscope on the right) Scale bar (left is 2 µm, right 5 µm)**

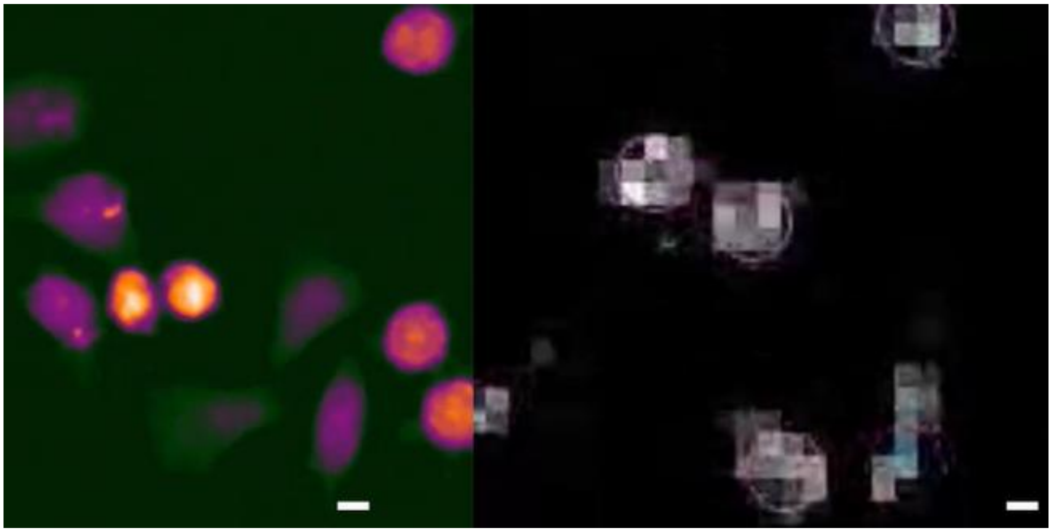

**Video C6 – Long term tracking of spheroid movement (L929 cultured in 1% vDMEM) using inverted LUT of defocus images**

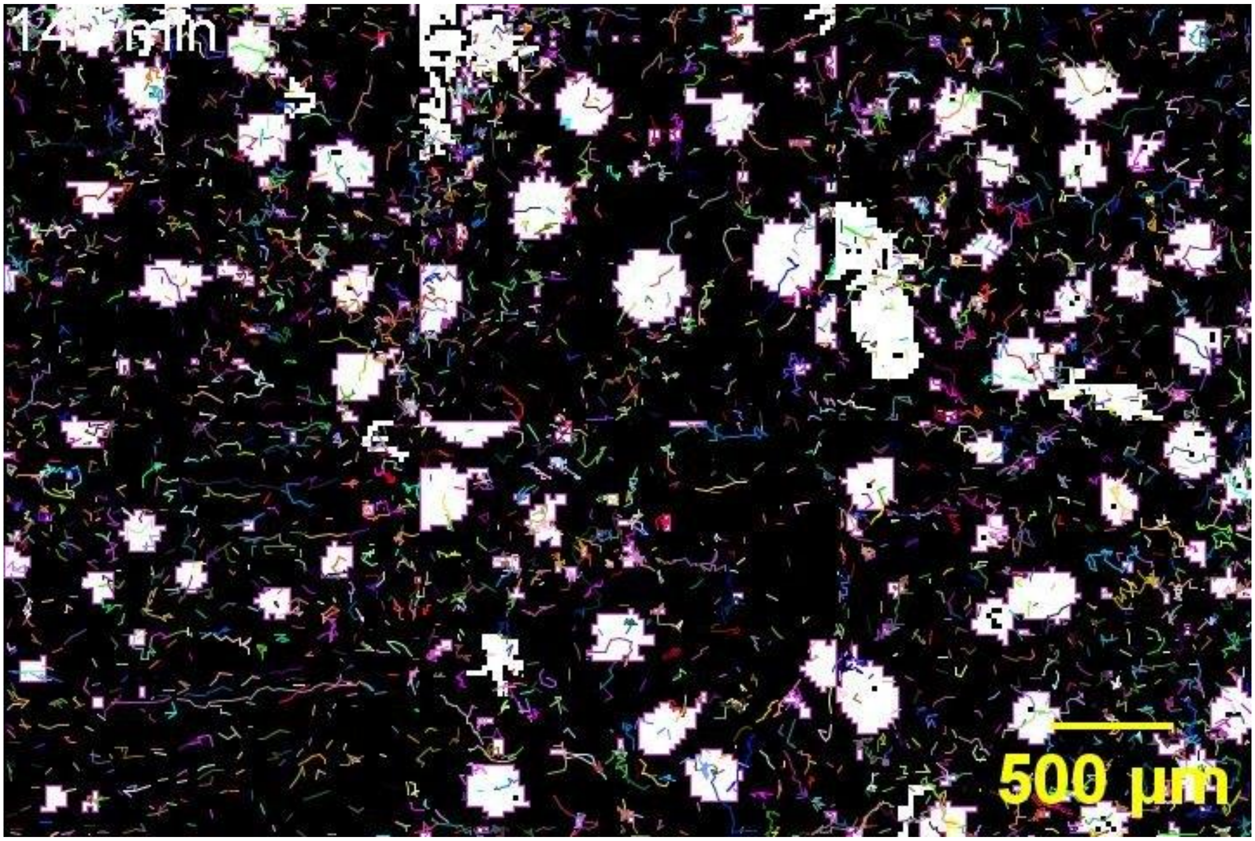
